# Neuropathological evidence of reduced amyloid beta and neurofibrillary tangles in multiple sclerosis cortex

**DOI:** 10.1101/2024.09.24.613639

**Authors:** J. Pansieri, M. Pisa, S. Yee, A. Gutnikova, E. Ridgeon, R. Hickman, J I Spencer, M. M. Esiri, G. C. DeLuca

**Affiliations:** Nuffield Department of Clinical Neurosciences, University of Oxford, Oxford, OX3 9DU, UK; Wessex Deanery, NHS England, Winchester, SO21 2RU, UK; Frimley Health NHS Foundation Trust, Frimley, Camberley GU16 7UJ; Foundation Medicine, Inc., 150 Second Street, Cambridge, MA 02141, USA; University College London Hospitals NHS Foundation Trust, 235 Euston Road, London, NW1 2BU, UK; Queen Square Institute of Neurology, University College London, London, WC1N 3BG, UK

**Author notes:** Corresponding author: Gabriele C. DeLuca, Nuffield Department of Clinical Neurosciences, Level 1, West Wing, John Radcliffe Hospital Oxford, United Kingdom, OX3 9DU.

## Abstract

Multiple sclerosis (MS) and Alzheimer’s disease (AD) are neurodegenerative diseases demonstrating age-related accumulation of disability. Inflammation lasting decades is a paradigmatic feature of MS pathology that variably relates to neurodegeneration, while the accumulation of Aβ plaques and neurofibrillary tangles (NFT) are cornerstones of AD pathology. However, few studies investigated the accumulation of amyloids in MS.

We investigated Aβ deposition and NFT density in temporal or frontal cortices derived from a large post-mortem cohort of MS (n=78) and age and sex-matched control (n=65) cases. We found reduced Aβ burden in MS cases compared with controls, particularly in cases below 65-years-of-age. NFT were similarly reduced in MS compared to controls, notably in cases above 65 years-of-age. Higher Aβ deposition predicted greater NFT density in MS. These findings suggest that MS-related factors may influence Aβ and NFT deposition and/or clearance. This work highlights new therapeutic perspectives relevant for both MS and AD.

## Introduction

Multiple sclerosis (MS) and Alzheimer’s disease (AD) are chronic neurodegenerative diseases, with both conditions showing an age-related progression of disability^1,2^. MS involves inflammation, demyelination, and neurodegeneration, leading to reduced quality of life. AD is primarily characterized by the accumulation of amyloid-beta (Aβ) and neurofibrillary tau tangles (NFTs), which are also present in areas commonly affected by MS^3,4^. However, how amyloid accumulation interacts with MS pathology, especially with aging, remains unclear.

Recent studies using PET imaging and plasma biomarkers indicate that Aβ burden may be lower in individuals with MS^5,6^. However, these methods may not differentiate between Aβ, and myelin or may be impacted by non-AD pathology, which limits the interpretation of the findings in the context of a condition like MS where cortical demyelination is prominent and co-morbidity is common^7^. Therefore, neuropathological validation is essential.

In the post-mortem study reported here, we show that Aβ deposition and NFT density are significantly lower in MS compared to controls, particularly in cortical demyelinated lesions. This supports the idea that MS-specific factors influence Aβ physiology, which has significant implications for understanding both AD and MS pathophysiology.

## Methods

### Study population

A post-mortem cohort of 141 individuals (aged 40 to 100 years) derived from non-dementia control (n=65) and pathologically-confirmed MS (n=76) cases was studied (Table 1). All autopsy material was obtained from the Oxford Brain Bank and UK MS Tissue Bank with relevant ethics committee approval (REC 15/SC/0639 and REC 08/MRE09/31+5, respectively), in accordance with the UK Human Tissue Act (2004).

**Table 1.** Demographics of MS and control cohort. (data are presented as mean, MS = multiple sclerosis ; N/A = not applicable ; M = male ; F = female ; PM = post-mortem, w/o = without).

|  | Control cases (n=65) | MS cases (n=76) | p values |
| --- | --- | --- | --- |
| Age at death (y) | 65.2 (range 40-100) | 64.0 (range 40-99) | 0.70 |
| Duration of disease (y) | N/A | 21.2 (range 1-62) | N/A |
| Sex | M = 39 ; F = 26 | M = 34 ; F = 42 | 0.09 |
| Brain weight (g) | 1381 (range 965-1760) | 1284 (range 1018-1794) | 0.002 (**) |
| PM interval (h) | 47.9 (range 8-160) | 48.9 (range 5-144) | 0.60 |
| Control and MS cohort used for neuronal assessments |  |  |  |
|  | Control cases (n=38) | MS cases (n=45) | p values |
| Number of cases (± median age, 64.6 yo) | below = 16 ; above = 22 | below = 21 ; above = 24 | N/A |
| Duration of disease (y) | N/A | 26.1 (range 1-62) | N/A |
| Sex | M = 22 ; F = 16 | M = 17 ; F = 28 | 0.08 |
| Brain weight (g) | 1356 (range 1017-1760) | 1270 (range 1018-1794) | 0.02 (*) |
| PM interval (h) | 50.7 (range 9-160) | 42.9 (range 5-92) | 0.70 |
| Frequency Aβ deposits | w/o Aβ = 17 ; with Aβ = 21 | w/o Aβ = 24 ; with Aβ = 21 | 0.51 |

### Immunohistochemistry on human post-mortem tissues

Immunohistochemistry was performed on formalin-fixed, paraffin-embedded blocks from the middle temporal gyrus of all cases. If no demyelinated lesions were found, the superior frontal gyrus was also sampled (n=7). Adjacent 6μm thick sections were stained for myelin (PLP), Aβ (4G8), and phosphorylated Tau (AT8), using chromogenic DAB and haematoxylin counterstaining (Supplementary Table 1, Supplementary Figure 1). Negative controls omitted primary and secondary antibodies.

### Demyelination Assessment

PLP-stained sections identified cortical MS lesions and non-lesional grey matter (NLGM), focusing on layers II-III, where Aβ deposition predominantly occurs.

### Aβ and Tau Deposition Assessment

Using digital scans at 400X magnification, 4G8 and AT8-stained sections were quantified in NLGM via a semi-automatic colour-based extraction method. Total Aβ and Tau expression was averaged across cortical layers, with a focused analysis in cortical layers II-III. Binary scores classified cases based on presence/absence of Aβ and Tau deposition.

In demyelinated cases, 174 lesions were identified and 156 lesion areas were delineated focusing on layers II and III for comparative analyses, and FOVs were sampled from core, border, peri-lesional, and NLGM regions. 138 lesions qualified given availability of lesion core, border, peri-lesion and NLGM areas involving cortical layers II and III.

### Neuronal density assessment

Neuronal densities were obtained by manually counting over 10,000 layer III pyramidal neurons in adjacent haematoxylin-stained lesional and NLGM FOVs. Neurons were identified by pyramidal/triangular shape with visible nucleus and nucleolus (Supplementary Figure 2).

To ensure consistent FOVs for each marker, microphotographs stained with each antibody and histochemical stain were co-aligned using Qupath software. Sections were anonymised to maintain blinding to disease status and 4G8 expression.

### Statistical analyses

Analyses were conducted using IBM SPSS® and GraphPad Prism by two independent researchers. Data for AT8 was normalised with log10 transformation. 4G8 and neuronal density data followed a normal distribution.

Cross-tab analysis evaluated differences in 4G8 expression and NFT densities (% of positive cases). Generalised linear models (GLM) compared mean 4G8 expression and AT8-positive neuron density between disease subgroups (MS vs. control, above/below median age-at-death), while generalised estimating equation models corrected for subject effects across cortical layers. GLM also assessed age and disease status effects on neuronal density, and their association with 4G8 expression.

Paired data for non-lesional vs. lesional areas were analysed using one-way ANOVA with Geisser-Greenhouse correction and Bonferroni’s multiple comparisons test. Mann-Whitney tests compared clinical features between MS and controls. Data are presented as mean ± SEM, with significance set at p<0.05. Bonferroni correction was applied for multiple comparisons.

## Results

### Demographics

Table 1 provides clinical details. MS cases and controls were matched for sex and age-at-death. The average disease duration for MS was 21.2 ± 15.2 years. The cohort was split into ‘younger’ and ‘older’ groups based on being below or above the median age-at-death (64.6 years), respectively. Brain weight was lower in MS cases compared to controls (p=0.002).

### Aβ is reduced in MS, especially at younger ages

Aβ was present in 44.6% of controls (29/65) and 35.5% of MS cases (27/76) (Figure 1A-B, p>0.1). In cortical layers II and III, Aβ expression tended to be lower in MS cases compared to controls (MS: 1.50 vs. controls: 2.48; Wald-X^2^ 3.31, p=0.069) after adjusting for age-at-death. Younger MS cases displayed a significant Aβ reduction compared to controls (MS: 0.39 vs. controls: 2.03; Wald-X^2^ 6.96, p=0.008), with no difference observed in older cases (Figure 1C-H), with similar results evaluating all cortical layers. Aβ associated with reduced brain weight in controls (p=0.047) but not in MS cases (data not shown). No link was found between Aβ expression and disease duration or sex.

**Figure 1.**
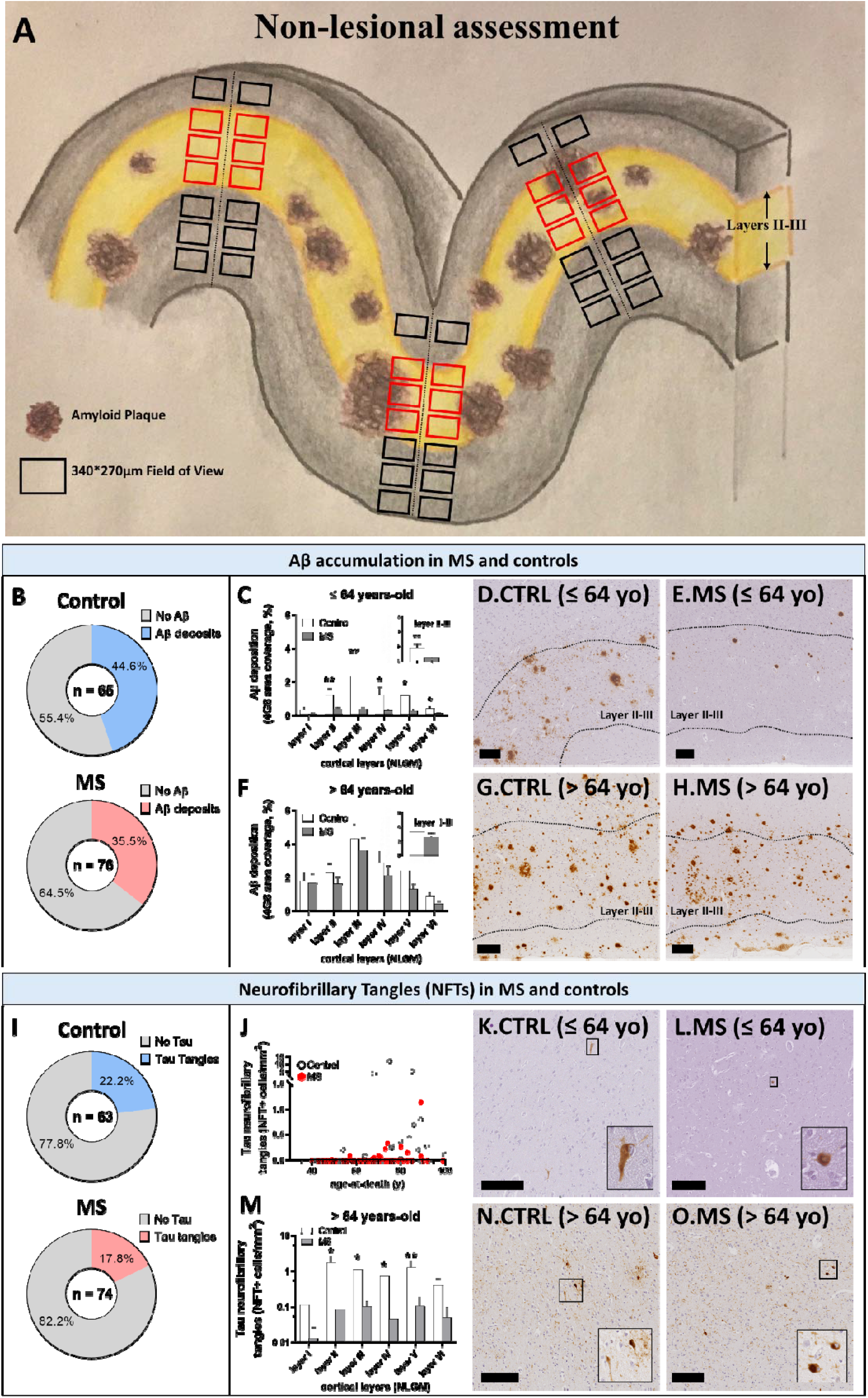
Amyloids are reduced in MS compared with control cortex. **(A)** 4G8 and AT8 expression was quantified in the non-lesional grey matter (NLGM) using pre-defined spaced trajectories (each 1 mm, when possible) perpendicular to the subpial surface of the cortex. For each trajectory, two fields of view (FOVs, black and red squares) of the same size were applied to NLGM layers, with the exception of layer III in which 4 FOVs were used due to the larger size of this layer, with a focused analysis on layer II-III (red squares). Using this method, more than 20,000 FOVs were quantified for each marker. **(B-H)** Aβ deposits in MS and controls. **(B)** A trend toward reduced number of MS cases compared with control showing Aβ deposition was observed. **(C-E)** Quantitation of 4G8 expression in cortical layers comparing NLGM from frontal and temporal cortices in control and MS cases shows a reduced Aβ deposition in younger than median age MS cases compared with controls. A particular attention on layer II-III (insert), the predominant site of Aβ deposition, show the same trend. **(F-H)** No difference in Aβ deposition comparing NLGM from frontal and temporal cortices was found comparing elder control and MS cases. **(I-O)** Neurofibrillary tangles (NFTs) density in MS and controls. **(I)** A trend toward reduced number of MS cases compared with control showing NFTs was observed. **(J)** NFTs were rarely found in young MS and control cases. **(K-O)** NFT density in cortical layers comparing NLGM from frontal and temporal cortices in control and MS cases shows a reduced NFT density in cortical layers in older than median age MS cases compared with controls. (generalised linear models; Data presented as mean□±□SEM ; * p < 0.05 ; MS = multiple sclerosis ; NLGM = non-lesional grey matter ; scale bar 200μm)

### NFTs are reduced in MS and predicted by Aβ expression

NFTs were found in 22.2% of controls (14/63) and 17.6% of MS cases (13/74) (Figure1I, p>0.1). NFT density was lower in MS compared to controls across all cortical layers (MS: 0.0131 vs. controls: 0.08; Wald 5.95, p=0.015), including layers III and V after adjusting for age.

In older cases, MS showed lower NFT density compared to controls (Figure 1J-O, NLGM: MS: 0.03 vs. controls: 0.13; Wald 5.24, p=0.022). No difference was observed in younger cases where NFT density was low in both MS and controls.

Higher NFT densities correlated with reduced brain weight in controls but not MS (data not shown). No association was found between NFT density and disease duration or sex. Aβ expression predicted NFT density in both MS and controls (data not shown, p < 0.0001).

### Aβ deposition is reduced in cortical lesions

Cortical lesions were detected in 76.3% of MS cases (58/76, Figure 2A-D). Of the 174 cortical lesions identified, 89.7% (156/174) affected cortical layers I-III, while 6.9% (12/174) affected layers IV-VI; 3.4% (6/174) affected WM. Of the total demyelinated areas, 52.2% affected layers II-III. Given the predominant amyloid deposition in layers II-III, quantitative measures of 4G8 and AT8 expression were assessed in demyelinated lesions affecting these layers. Controls had no demyelination.

**Figure 2.**
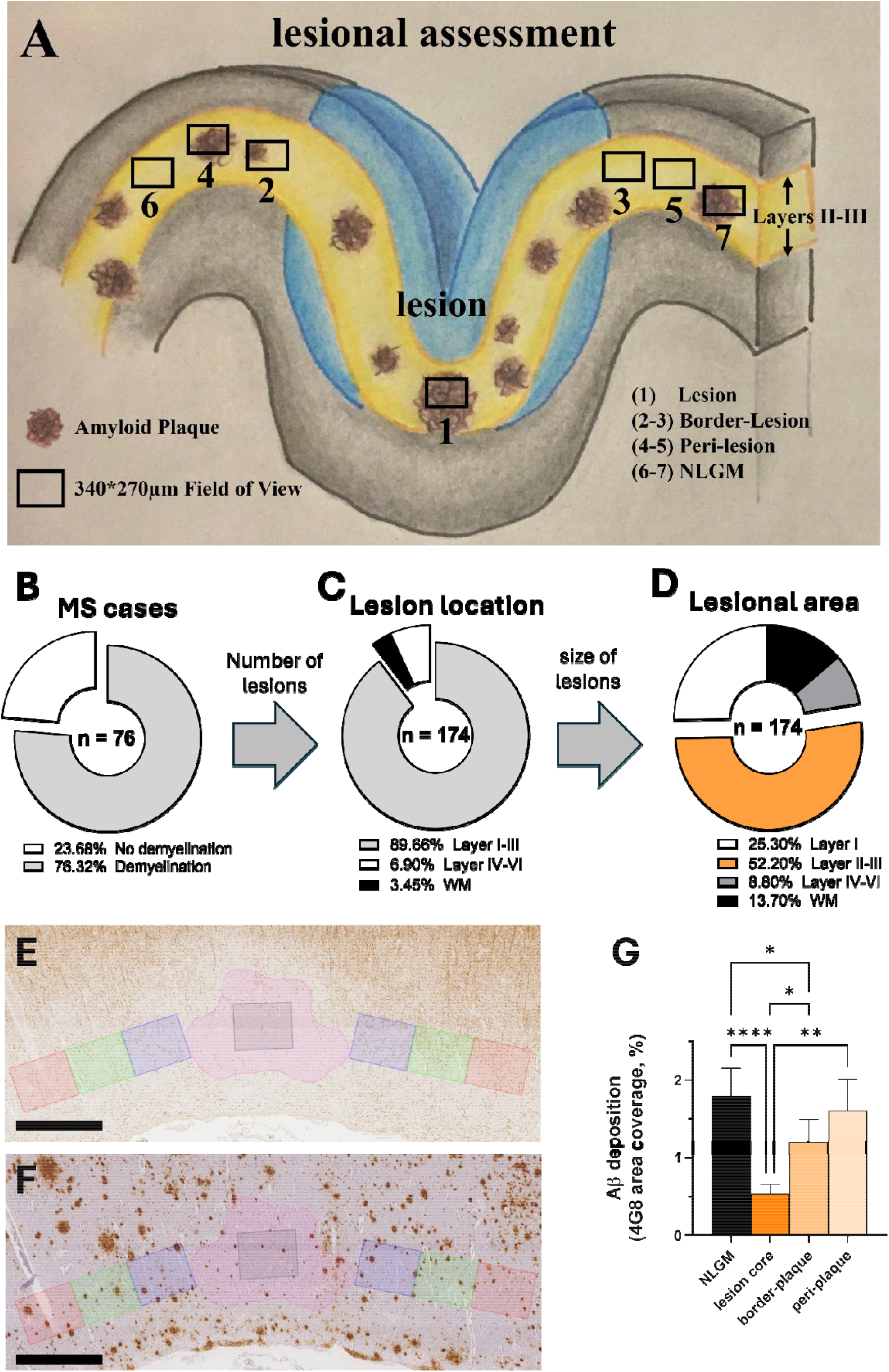
Aβ burden and demyelination in MS cortex. **(A)** MS lesions were defined by complete loss of myelin. In cases with demyelination, the entire lesional area was delineated for each lesion and a FOV in the core of the lesion was used (black square 1). When available, two additional FOVs of the same size matched for cortical layer location on each side of the lesion were used for border, peri-lesional, and non-lesional areas as follows: border region – from lesion edge up to 340 μm of adjacent NLGM (black squares 2-3); peri-lesional region –340 to 680 μm from lesion edge (black squares 4-5); NLGM region more than 680 μm from lesion edge (black squares 6-7). Given the preferential deposition of Aβ in layers II and III, we focused our comparative analyses of 4G8 expression in and outside of lesions affecting theses cortical layers. **(B)** Proportion of MS cases (n=76) showing demyelination. **(C)** Proportion of demyelinated lesions (n=174) depending on their location in the grey matter layers and white matter. **(D)** Distribution of lesional area size on the total demyelinated area in white matter and grey matter observed in our MS cohort. **(E-G)** Representative field of view in a demyelinated lesion identified by **(E)** PLP staining and **(F)** corresponding area with 4G8 staining. Pink field of view (FOV) represent the total demyelinated plaque area, while squared FOVs represent the lesion core (light grey), border-plaque area (light blue), peri-plaque area (light green) and non-lesional grey matter (light red) in corresponding cortical layers. **(G)** Quantitation of Aβ deposition (4G8 expression) comparing demyelinated lesions affecting the layer II-III and corresponding non-lesional grey matter layers (n=156) shows a reduced Aβ deposition in lesional and border-plaque areas (ANOVA and post-hoc paired t-test ; Data presented as mean□±□SEM ; * p < 0.05 ; ** p < 0.01 ; **** p<0.0001 ; MS = multiple sclerosis ; NLGM = non-lesional grey matter; scale bar 500 μm)

Aβ deposition was lower within the lesion core compared to border, perilesional, and NLGM areas (Figure 2E-G). No difference in NFT density within and outside lesions was found (data not shown). Age and sex did not impact Aβ expression and NFT density in cortical lesions.

### Neuronal density

No differences were observed in neuronal density between MS and control (Supplementary Figure 3). However, neuronal densities were reduced in MS cases with Aβ deposits compared to MS cases without Aβ deposits, a finding driven by younger cases (n=21; 51.29 ± 16.23 vs 72.98 ± 22.34, p=0.045).

No association between Aβ deposition and neuronal counts was found in controls.

## Discussion

Our study offers a detailed analysis of Aβ and tau accumulation in post-mortem MS and non-neurological control brains across the lifespan. Key findings include: (1) Aβ and NFT accumulation are reduced in MS non-lesional cortex,(2) Aβ deposition is notably lower in demyelinated lesions than in non-lesional areas in MS; (3) Aβ deposition is associated with lower neuronal densities in younger MS cases. These results suggest complex interactions between age, amyloid accumulation, neuronal loss, and inflammation, with implications on our understanding of cognitive impairment determinants in MS and neurodegenerative diseases like AD.

Previous MS studies focused on amyloid protein intermediates, biomarkers, or animal models, such as experimental autoimmune encephalomyelitis (EAE)^8–10^. Our finding of reduced Aβ expression in MS aligns with recent PET imaging and biomarker studies but provides more detailed, quantitative neuropathological insights^5^. A previous post-mortem study did not find similar reductions in MS^11^, possibly due to differences in cohort size and protocol. Our study’s focus on amyloid-rich cortical layers^12,13^ and larger cohort size strengthens our conclusions.

The mechanisms behind reduced amyloids in MS remain unclear. Inflammatory insults in animal models where amyloids are reduced^14^ suggest chronic inflammation in MS may influence amyloid deposition or clearance. Altered blood-brain barrier function in MS may also play a role. MS therapies, not yet available when these patients died, do not explain our findings. Interestingly, amyloid administration reduces EAE severity^15,16^, suggesting amyloids might help maintain immune homeostasis in MS. Future research should explore relationships between amyloids and inflammation in well characterised post-mortem MS brain tissue.

Though autopsy series often involve severe cases, our large cohort matched for age-at-death across the lifespan mitigates this bias. Lack of cognitive data limits clinical correlation, and factors like fixation time impacted the exploration of inflammatory markers. However, our Aβ findings, confirmed via silver stain (data not shown), and consistent reductions in Aβ and Tau, particularly within lesions of the same case, offer robust internal controls.

This study reveals reduced amyloid accumulation in MS brains, crucial for understanding MS and other neurological diseases, and highlights an overlooked interaction between Aβ and Tau with MS pathology.

## Acknowledgements

This study was funded by the Biomedical Research Centre, Oxford, National Institute of Health Research, the Oxford-Quinnipiac-Trinity Health partnership, the UK MS Society, and Oxford University Clinical Academic Graduate School. We would like to thank Dr Gargi Banerjee for their contributions.

