## Supplemental Files for "Neuropathological evidence of reduced amyloid beta and neurofibrillary tangles in multiple sclerosis cortex"

**Running head: Amyloid reduction in multiple sclerosis cortex**

J. Pansieri^1^ PhD, M. Pisa^1^ MD, S. Yee^1^ DPhil, A. Gutnikova^2^ MA (Oxon) BMBCh MRCPCH, E. Ridgeon^3^ BA BMBCh MSc FRCA, R. Hickman^4^ MD MRCS, J I Spencer^5,6^ BMBCh, M. M. Esiri^1^ DM FRCPath, G. C. DeLuca^1*^ MD, DPhil, FRCPath

*^1^ Nuffield Department of Clinical Neurosciences, University of Oxford, Oxford, OX3 9DU, UK*

*^2^Wessex Deanery, NHS England, Winchester, SO21 2RU, UK*

*^3^Frimley Health NHS Foundation Trust, Frimley, Camberley GU16 7UJ*

*^4^Foundation Medicine, Inc., 150 Second Street, Cambridge, MA 02141, USA*

*^5^University College London Hospitals NHS Foundation Trust, 235 Euston Road, London, NW1 2BU, UK*

*^6^Queen Square Institute of Neurology, University College London, London, WC1N 3BG, UK.*

* Corresponding author:

Gabriele C. DeLuca

Nuffield Department of Clinical Neurosciences

Level 1, West Wing, John Radcliffe Hospital

Oxford, United Kingdom, OX3 9DU

| **Target** | **Primary Antibody** | **Antibody dilution** | **Clone** | **Antigen Retrieval** | **Incubation Settings** |
| --- | --- | --- | --- | --- | --- |
| PLP | Biorad #MCA839G | 1 : 1 000 | monoclonal | Citrate pH6 Microwave | 1h RT |
| 4G8 | Biolegend  #SIG-39220 | 1 : 24 000 | monoclonal | Formic acid | ON RT |
| AT8 | Innogenetics | 1 : 1 500 | monoclonal | None | 1h RT |

**Supplementary Table 1. Antibodies used in staining procedures presented in this article.** (ON = overnight ; RT = room temperature)

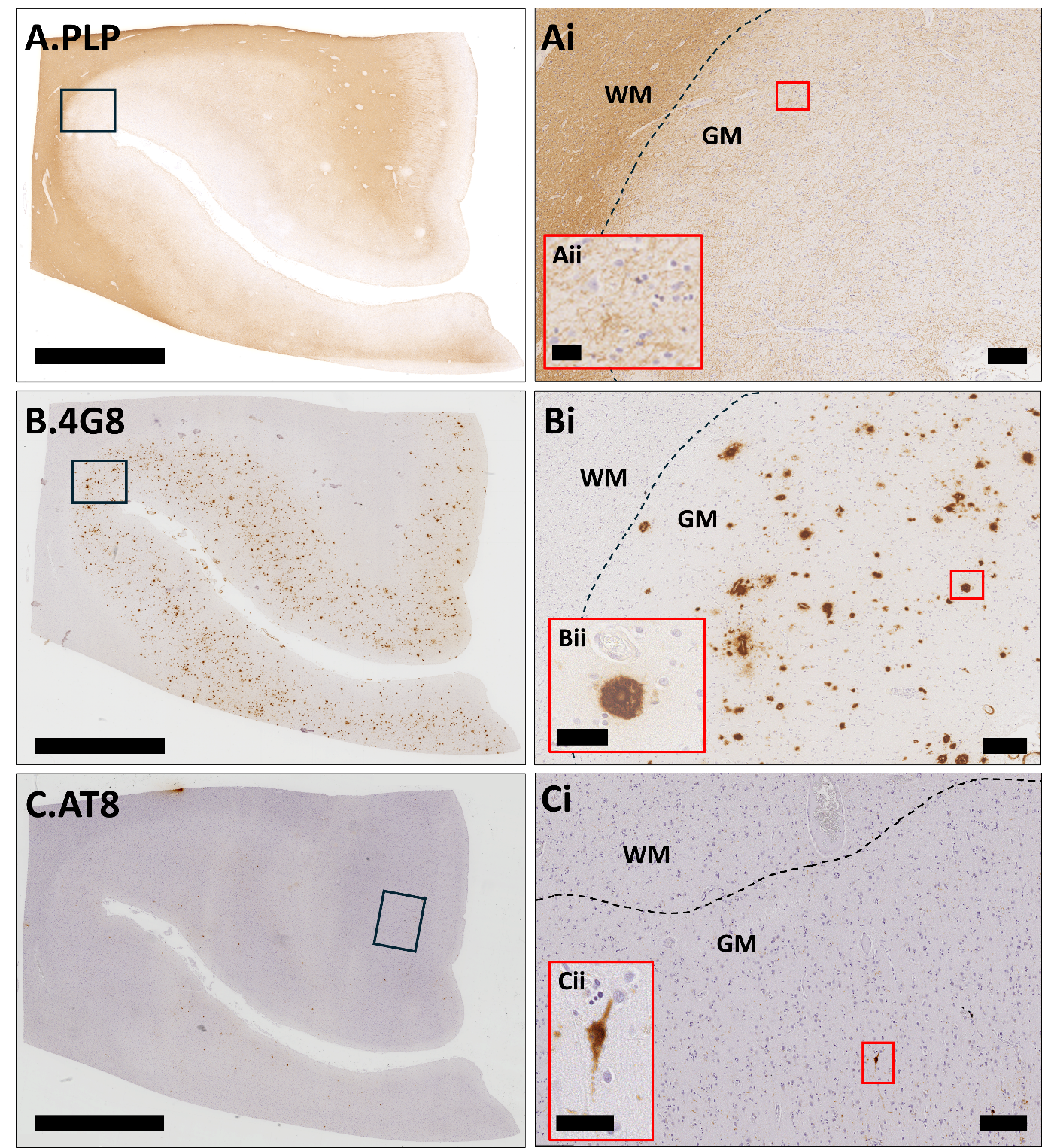

**Supplementary Figure 1. Representative immunohistochemistry.** Adjacent sections of a MS case were immunostained for **(A)** PLP, **(B)** 4G8 and **(C)** AT8 (scale bar 5mm). Higher magnification of each marker (black squares) is represented in Ai, Bi and Ci, respectively (scale bar 200µm). Higher magnification (red squares) for myelin, Aβ deposits, Tau accumulation (with neurofibrillary tangle in the insert) are presented in Aii, Bii, and Cii, respectively (scale bar 50µm). (WM = white matter ; GM = grey matter ; dotted line shows WM/GM boundaries).

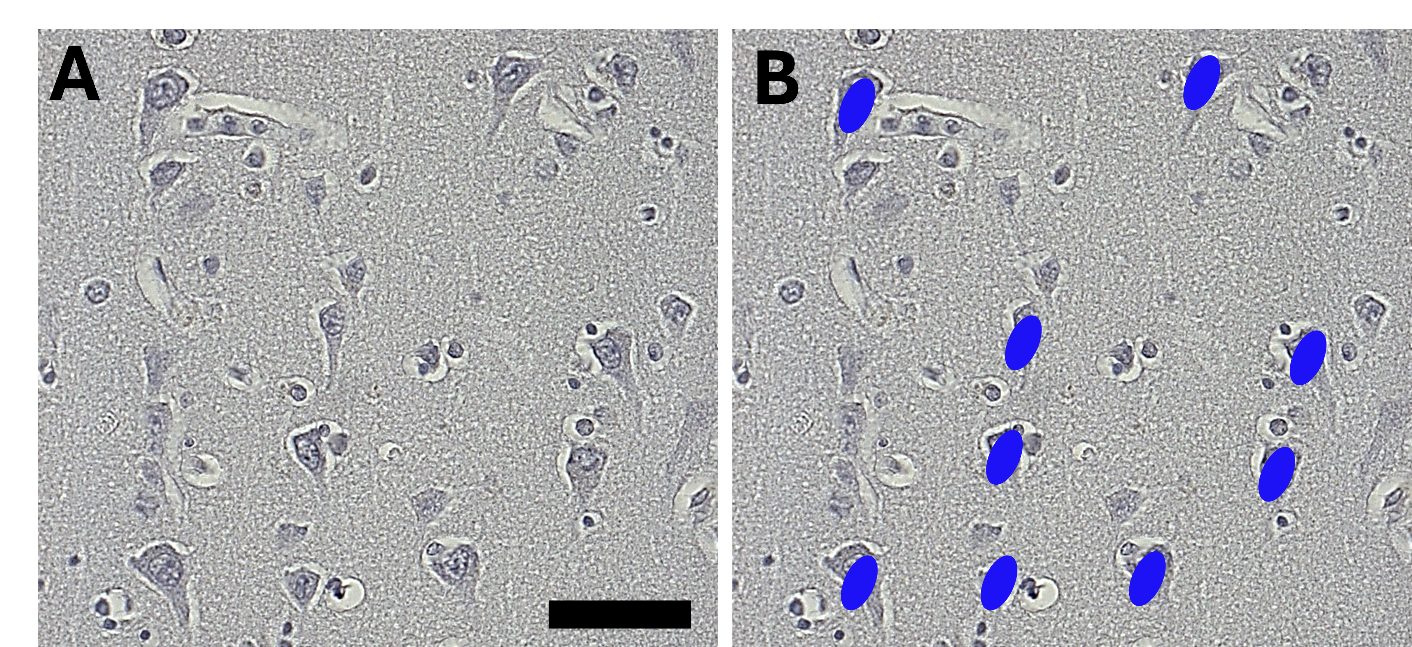

**Supplementary Figure 2. Neuronal counting methods.(A)** A sample image in cortical layer 3 is shown. **(B)** The same image is presented with counting markers. Blue circles indicate the pyramidal neurons counted for neuronal densities, where only neurons showing a pyramidal shape with visible nucleolus were taken into account. (Scale bar 50 µm).

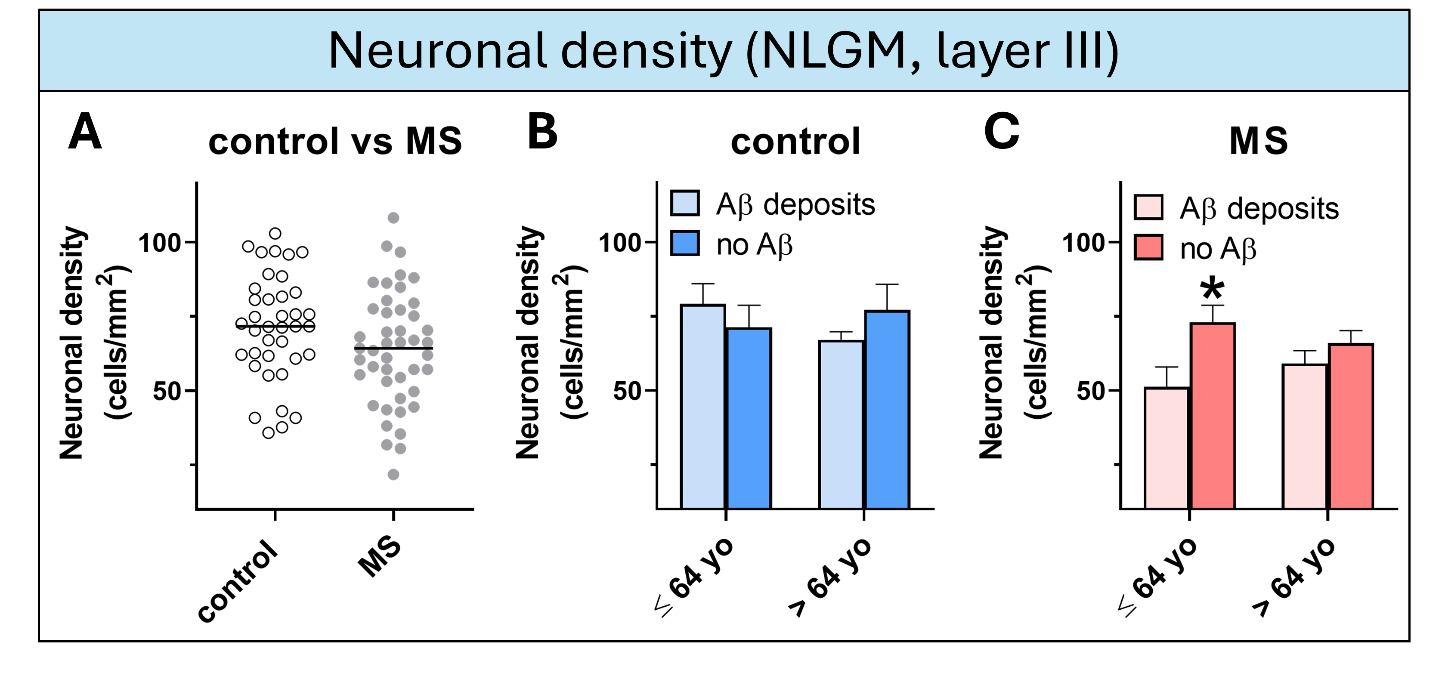

**Supplementary Figure 3.** **Neuronal density and Aβ deposition in MS compared with control cases**. **(****A)** No significant differences were found in neuronal densities comparing control and MS cases. **(B-C)** In MS cases younger than median age, an increase in layer III neuronal density was found in cases without Aβ deposition compared with cases with Aβ deposition, while no difference was observed in control cases. (data presented as mean ± SEM ; * p < 0.05 ; ** p < 0.01 ; MS = multiple sclerosis ; NLGM = non-lesional grey matter)
